# Hepatic Acetate is an Essential Fuel for the Heart during Energy-Deprived States

**DOI:** 10.64898/2026.05.29.728677

**Authors:** Jinyang Wang, Tong Ren, Yiran Zhang, Lizheng Yao, Hao Yu, Ziyuan Wang, Ying Zhang, Songyang Li, Shitao Liang, Jia Li, Bin Jiang, Jiahuai Han, Gang Liu, Qinxi Li

## Abstract

Energy supply is fundamental to cardiac performance, yet the mechanisms by which the heart adapts to nutrient scarcity remain incompletely understood. Here, we identify acetate as a pivotal metabolic substrate that sustains cardiac contractile function during fasting-induced energy deficiency. Fasted mice reveal significant elevation of circulating acetate derived from hepatic fatty acid catabolism. Physiological concentrations of acetate alone can maintain *ex vivo* beating and electrical stability in Langendorff-perfused hearts. Primary mouse cardiomyocytes preferentially utilize acetate under energy-restricted conditions, which is abolished by knockdown of mitochondrial Acss1, the enzyme responsible for converting acetate to acetyl-CoA. Isotopic tracing with ^13^C-acetate demonstrated selective cardiac uptake during fasting. *In vivo*, acetate supplementation preserved heart rate and systolic function in fasted wild-type mice but not in Acss1-deficient hearts. These findings reveal an Acss1-dependent metabolic pathway enabling the heart to harness hepatic acetate as an endogenous fuel, representing a nutrient stress-responsive adaptation based on inter-organ crosstalk and implying therapeutic potential of acetate to energy-deficient heart diseases.

## INTRODUCTION

The heart, as the central organ of the circulatory system, demands a continuous and efficient energy supply to sustain unremitting contractions. To meet this exceptionally high energy demand, cardiomyocytes primarily rely on tightly regulated metabolism of fatty acids and glucose ^1–3^. However, under physiologically stressful conditions such as fasting, malnutrition or ischemia, systemic energy scarcity poses significant challenges to cardiac energetic homeostasis ^4–7^. Despite these constraints, the heart exhibits remarkable functional stability, suggesting the existence of intrinsic adaptive mechanisms. Elucidating how the heart modulates substrate preference, metabolic flux, and energy partitioning under energy stress is critical for understanding cardiac resilience and may inform therapeutic strategies for metabolic cardiomyopathies and energy-deficient heart diseases.

## RESULTS

### 1. Acetate Supports Sustained Cardiac Beating as a Sole Energy Substrate *ex vivo*

While the heart can oxidize various substrates—including glucose, pyruvate, lactate, fatty acids, and ketone bodies ^1,8,9^—how it dynamically adapts substrate preference during fasting remains poorly defined. To further characterize fasting-induced metabolic changes, we analyzed various energy substances in serum of mice fasted for 18 hours employing HP-CIL LC-MS. As we reported previously ^10^, the levels of circulating acetate (the fatty acid with shortest chain) were dramatically risen from 166.1 ± 37.7 μM in normal state to 454.2 ± 92.8 μM in response to fasting (Figure 1A). Other short-chain fatty acids exhibited different alterations, with propanoate being increased and butyrate keeping no change (Figure 1B-C). Long-chain fatty acids, especially C16 and C18 fatty acids (the most abundant side chains in triacylglycerol), as well as C14, C20, C22 and C24 fatty acids which also participate frequently the construction of triacylglycerol, were markedly elevated, indicating the mobilization of triacylglycerol in adipose tissue (Figure S1A-B). Expectedly, ketone bodies 3-hydroxybutyrate (3-HB) and acetoacetate (AcAc) were significantly upregulated (Figure 1D-E). These results suggest that, during starvation, triacylglycerols are catabolized to release free fatty acids, which are subsequently converted in the liver into acetate and ketone bodies. In contrast, the levels of glucose and glucose-related metabolites such as lactate and ribose were markedly decreased, demonstrating the lack of glucose and its downstream metabolites under fasting (Figure S1C-E). As to amino acids, the majority of amino acids tested remained unchanged, which suggest that a 18-hour fasting is not as severe as to induce the broad degradation of proteins of peripheral tissues such as muscle (Figure S1F).

**Figure 1.**
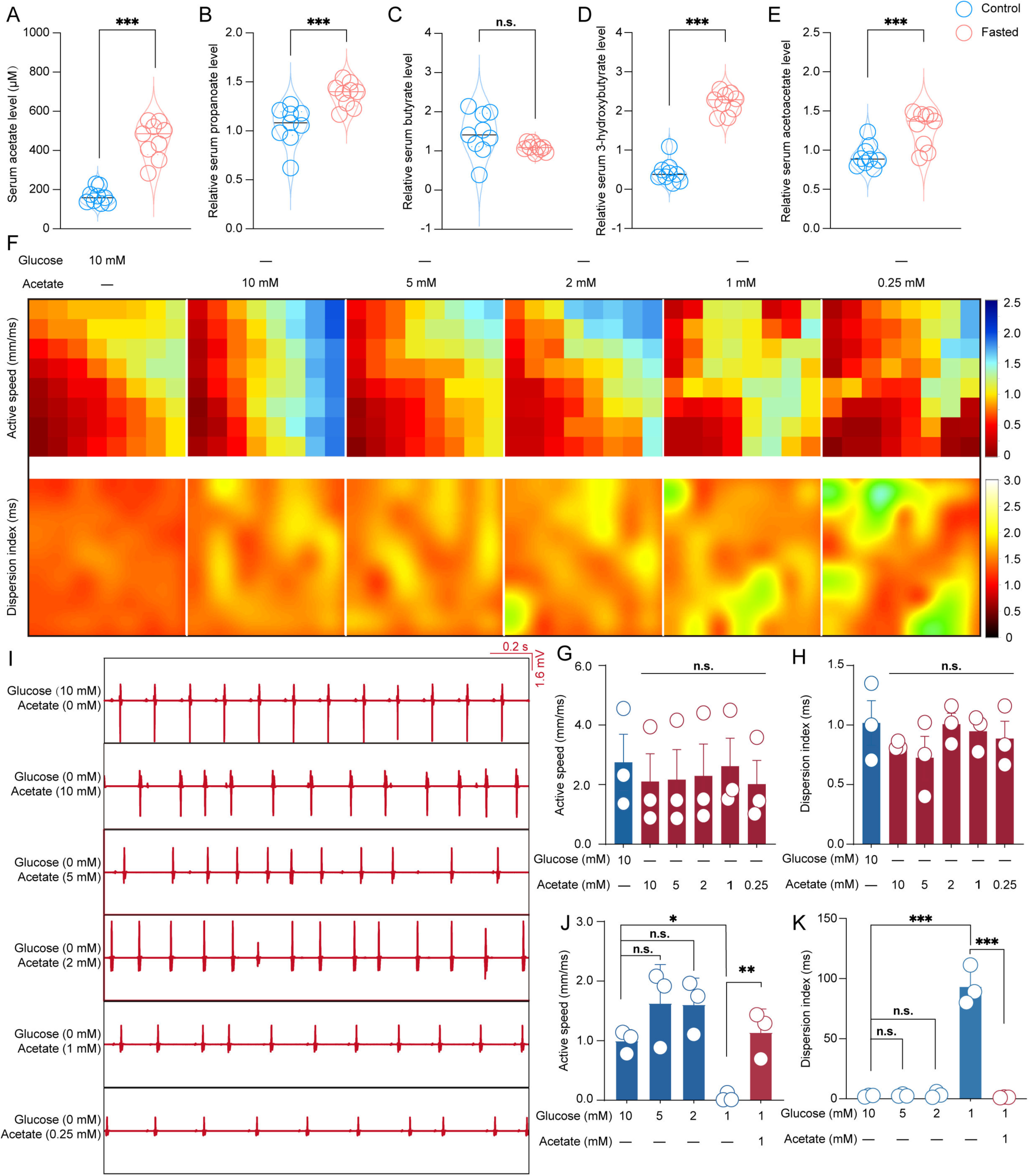
Acetate Supports Sustained Cardiac Beating as a Sole Energy Substrate ex vivo. (A-E) The serum levels of acetate (A), propanoate (B), butyrate (C), 3-hydroxybutyrate (D) and acetoacetate (E) in mice fasted for 18 h (n = 9). (F-K) Representative maps of ventricular activation speed and dispersion index (F), quantification of activation speed (G, J) and dispersion index (H, K), and representative electrocardiogram traces (I) from *ex vivo* Langendorff-perfused wild-type mouse hearts under indicated perfusion conditions (n = 3). Data are presented as mean ± standard deviation (SD) from ≥3 biological replicates. For (A-E), two-tailed unpaired Student’s *t*-tests were used. For (G, H, J and K), statistical significance was determined by one-way ANOVA comparing each treatment group (\**p* < 0.05, \*\**p* < 0.01, \*\*\**p* < 0.001, n.s., no significant difference).

To evaluate whether fasting-induced metabolites can sustain cardiac function, we conducted *ex vivo* perfusion of Langendorff hearts from wild-type mice ^11^. Individual metabolites were added to a carbon-free perfusate, and myocardial electrical activity was monitored using a 64-channel epicardial multielectrode array. At 10 mM, acetate, glucose, lactate, pyruvate and butyrate, but not propanoate, 3-HB and AcAc, separately supported continuous myocardial contractions with preserved conduction dispersion and sinus rhythm (Figure 1F-I and S2). Notably, palmitate—the most abundant circulating free fatty acid in serum —also failed to support sustained myocardial contraction around its physiological concentrations (0.5 mM) (Figure S2A-C) ^12^. Among the tested metabolites, acetate was the only one both significantly elevated in fasting serum and capable of supporting continuous myocardial contraction as a sole carbon source. To further assess its physiological relevance, we titrated acetate concentrations and found that at near-physiological levels (250 μM) ^13^, acetate alone still could sustain preserved conduction dispersion and stable cardiac beating in the *ex vivo* Langendorff model (Figure 1F-I). Moreover, acetate supplementation rescued the loss of sustained myocardial contraction induced by reduced glucose availability (Figure 1J-K and S3). These findings identify acetate as a key metabolic substrate that supports cardiac function during fasting-induced energy restriction.

### 2. Acetate Sustains Cardiomyocyte Beating by Preserving Electrical Activity

To investigate how acetate sustains cardiac contractions, we assessed its utilization across multiple primary murine cell types. Notably, only cardiomyocytes (CMs) exhibited substantial ^13^C-acetate consumption under glucose-deprived conditions (Figure 2A). Moreover, among various substrates, acetate was preferentially utilized by CMs when glucose was absent (Figure 2B). Under energy-restricted conditions (glucose free plus inhibition of mitochondrial carnitine-acylcarnitine translocase with Etomoxir)—where both glucose and fatty acid availability were limited—spontaneous CM beating was markedly reduced, but was significantly restored by acetate supplementation (Figure 2C-D).

**Figure 2.**
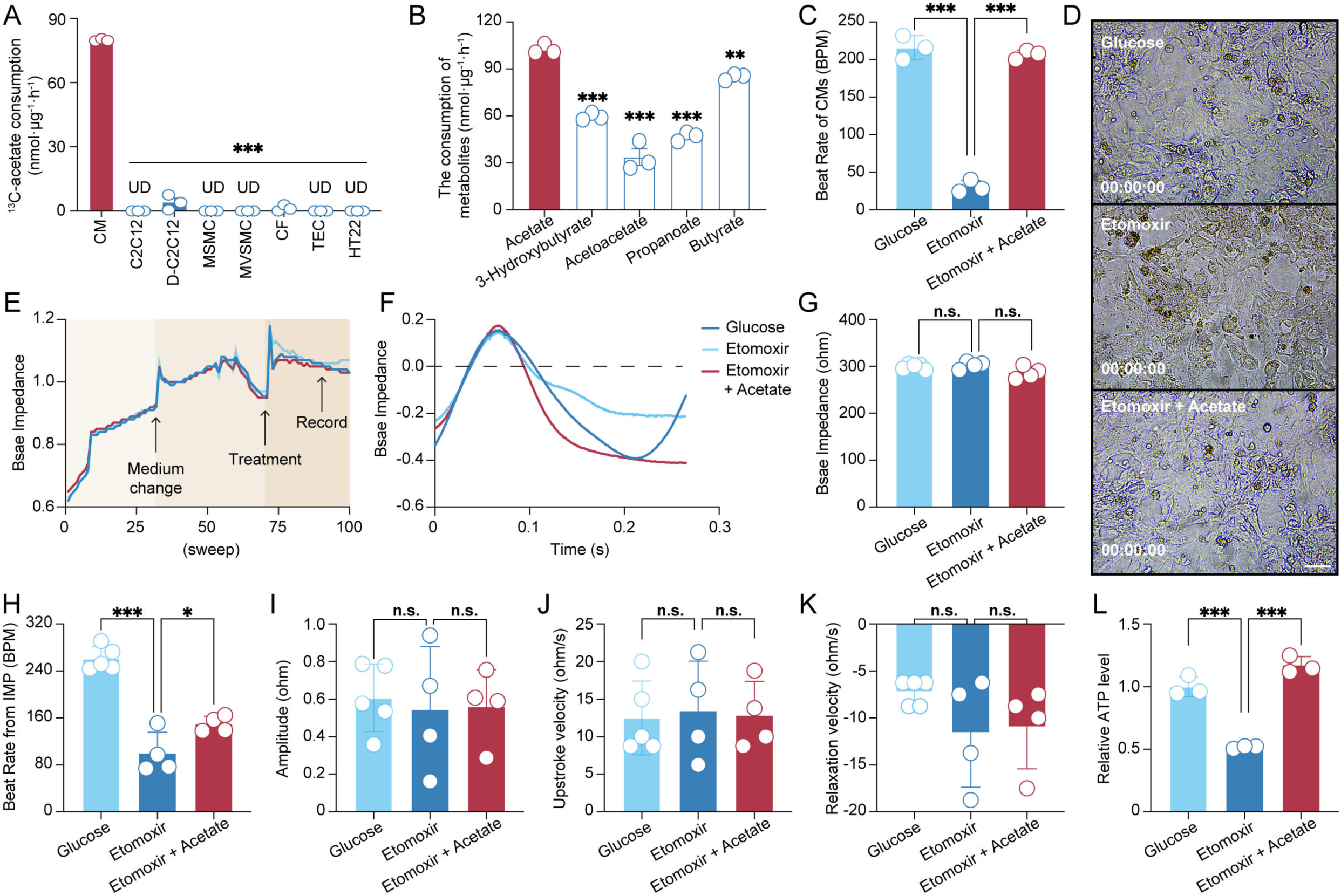
Acetate Sustains Cardiomyocyte Beating by Preserving Electrical Activity. (A) Consumption of 2-^13^C-acetate by the indicated cell types cultured in 2-^13^C-acetate-containing medium for 24 h (n = 3). Statistical comparisons were performed between each cell type and cardiomyocytes (CMs). (B) Consumption of individual metabolites by CMs cultured for 24 h in media containing the corresponding single metabolite (n = 3). Statistical comparisons were performed between each metabolite and acetate. (C and D) Representative traces and quantification of spontaneous beating of CMs cultured for 6 h under energy-restricted conditions (glucose-free medium supplemented with 40 μM etomoxir) with or without acetate supplementation. Scale bars represent 40 μm. Video data (D) are provided in the Supplementary Materials. (E-K) Impedance-based analysis of CMs under the same energy-restricted conditions as in (C), showing representative base impedance recordings over time (E), baseline impedance (F and G), beat rate (H), contraction amplitude (I), upstroke velocity (J), and relaxation velocity (K) after 120 h of pacing (n = 4). (L) Intracellular ATP levels in CMs cultured for 6 h under energy-restricted conditions with acetate supplementation (n = 3). Values are expressed as mean ± SD from ≥ 3 biological replicates and analyzed using one-way ANOVA (\**p* < 0.05, \*\**p* < 0.01, \*\*\**p* < 0.001, n.s., no significant difference). Abbreviations: CM, cardiomyocyte; D-C2C12, differentiated C2C12 myotubes; MSMC, primary mouse skeletal muscle cell; MVSMC, primary mouse vascular smooth muscle cell; CF, cardiac fibroblast; TEC, primary mouse renal proximal tubular epithelial cell; UD, undetectable; BMP, beats per minute.

To further evaluate this effect, CM electrical activity was assessed using the impedance (IMP) mode of the CardioExcyte 96 system ^14^. Impedance recordings revealed stable and rhythmic fluctuations corresponding to regular contractile activity across experimental conditions (Figure 2E-G). Under nutrient-restricted conditions, the beat rate was markedly reduced, whereas acetate supplementation restored beating frequency (Figure 2H). In contrast, parameters reflecting contractile dynamics—including amplitude (indicative of contraction strength), upstroke velocity (reflecting contraction kinetics), and relaxation velocity (reflecting relaxation kinetics)—remained largely unchanged across conditions (Figure 2I-K). Consistent with its role as an energetic substrate, acetate also restored intracellular ATP level in nutrient-deprived CMs (Figure 2L). In contrast, acetate exerted minimal effects on the beating rate and energetic index under nutrient-rich conditions (Figure S4). Together, these findings indicate that acetate supports CMs contractile function under energy stress by sustaining ATP level and electrical excitability.

### 3. Acss1-Mediated Acetate Utilization Sustains Cardiac Contraction under Energy Deficiency

Acetate metabolism is mediated by two acyl-CoA synthetase short-chain family isoforms: mitochondrial Acss1 which synthesizes acetyl-CoA from acetate for degradative oxidation, and cytosolic Acss2 which converts acetate to acetyl-CoA for lipid biosynthesis ^15^. Mouse cardiac tissue exhibited selective enrichment of Acss1, with negligible Acss2 expression as detected by Western blot (Figure 3A). Genetic silencing of Acss1 in mouse primary CMs efficiently reduced the acetate utilization (Figure 3B). Consistently, the loss of Acss1 abrogated the capacity of acetate to rescue the beating rate of CMs during nutrient deprivation (Figure S5). Extending to bioenergetic metrics, acetate-endowed ATP restoration upon nutrient deprivation was abolished by Acss1 depletion (Figure 3C). We next sought to investigate the functional requirement of Acss1 in whole hearts using an *ex vivo* Langendorff perfusion system. Strikingly, the absence of Acss1 rendered acetate completely ineffective in supporting myocardial contractions (Figure 3D-G and S6A). Taken together, acetate relies on Acss1 to bolster cardiac contractile function under energy-deprived conditions by sustaining ATP supply.

**Figure 3.**
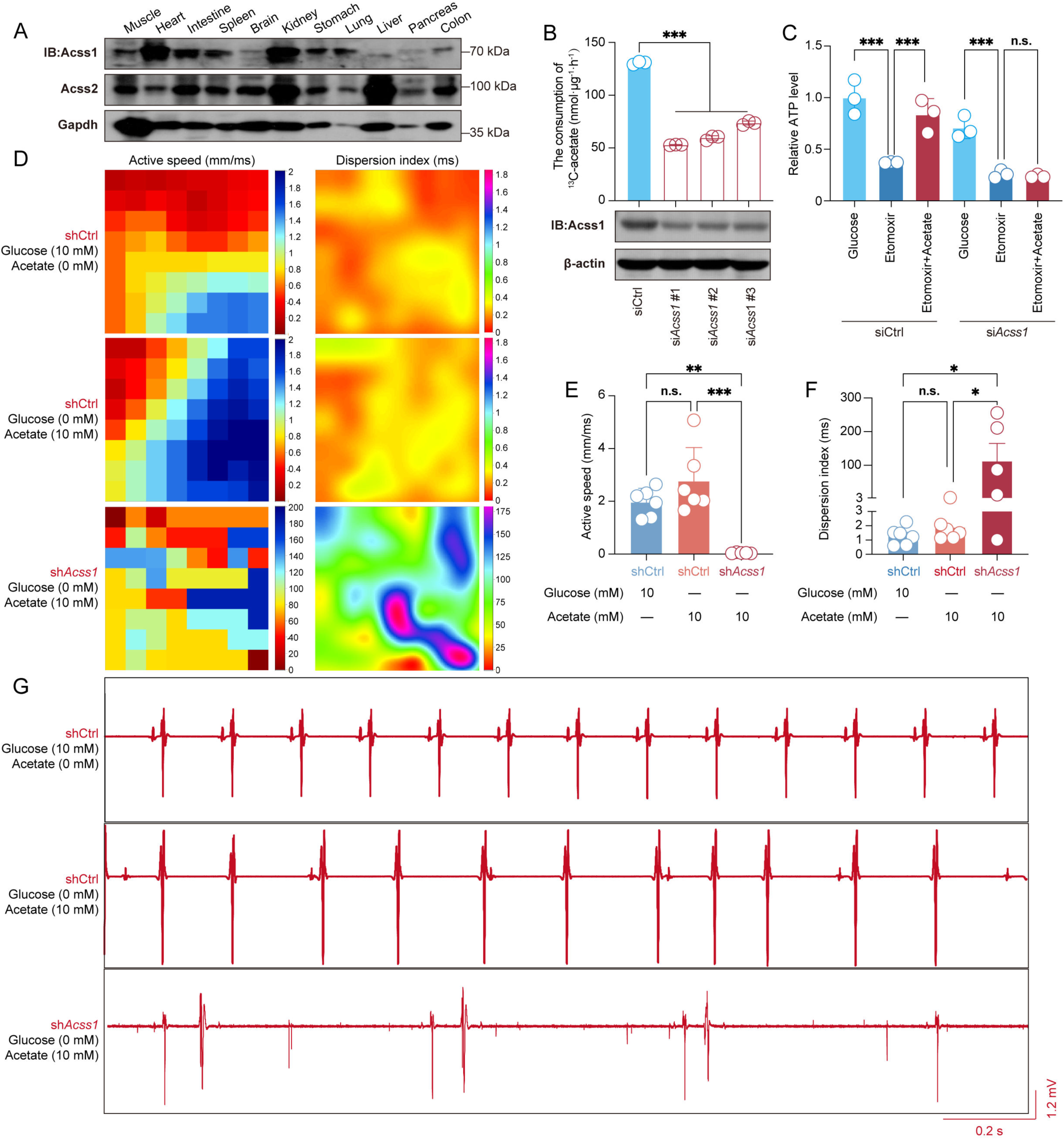
Acss1-Mediated Acetate Utilization Sustains Cardiac Contraction under Energy Deficiency. (A) Protein levels of Acss1 and Acss2 in multiple tissues from C57BL/6 mice. (B) 2-^13^C-acetate consumption by CMs with RNAi-mediated *Acss1* knockdown after 24 h (n = 3). (C) Intracellular ATP levels in *Acss1*-silenced CMs cultured under energy-restriction conditions with acetate supplementation (n = 3). (D-G) Representative ventricular activation speed and dispersion index (D), quantification of activation speed (E) and dispersion index (F), and electrocardiogram traces (G) from Langendorff-perfused mouse hearts with AAV9-mediated cardiac-specific *Acss1* knockdown exposed to acetate (n = 6). Values are expressed as mean ± SD from ≥ 3 biological replicates and analyzed using one-way ANOVA (B, E and F) or two-way ANOVA (C) (\**p* < 0.05, \*\**p* < 0.01, \*\*\**p* < 0.001, n.s., no significant difference).

### 4. Hepatic Acetate Maintains Cardiac Function via Acss1 *in vivo* under Energy Restriction

We next investigated the *in vivo* relevance of circulating acetate during energy deprivation. Building on our previous findings that hepatic fatty acid oxidation is a major source of circulating acetate during fasting ^10^, we conducted *in vivo* isotopic tracing with 2-^13^C-acetate in fasted mice and performed LC-MS-based metabolite analysis. The heart exhibited significantly higher ^13^C-acetate incorporation compared to other organs, indicating a preferential utilization of circulating acetate in cardiac tissue (Figure 4A).

**Figure 4.**
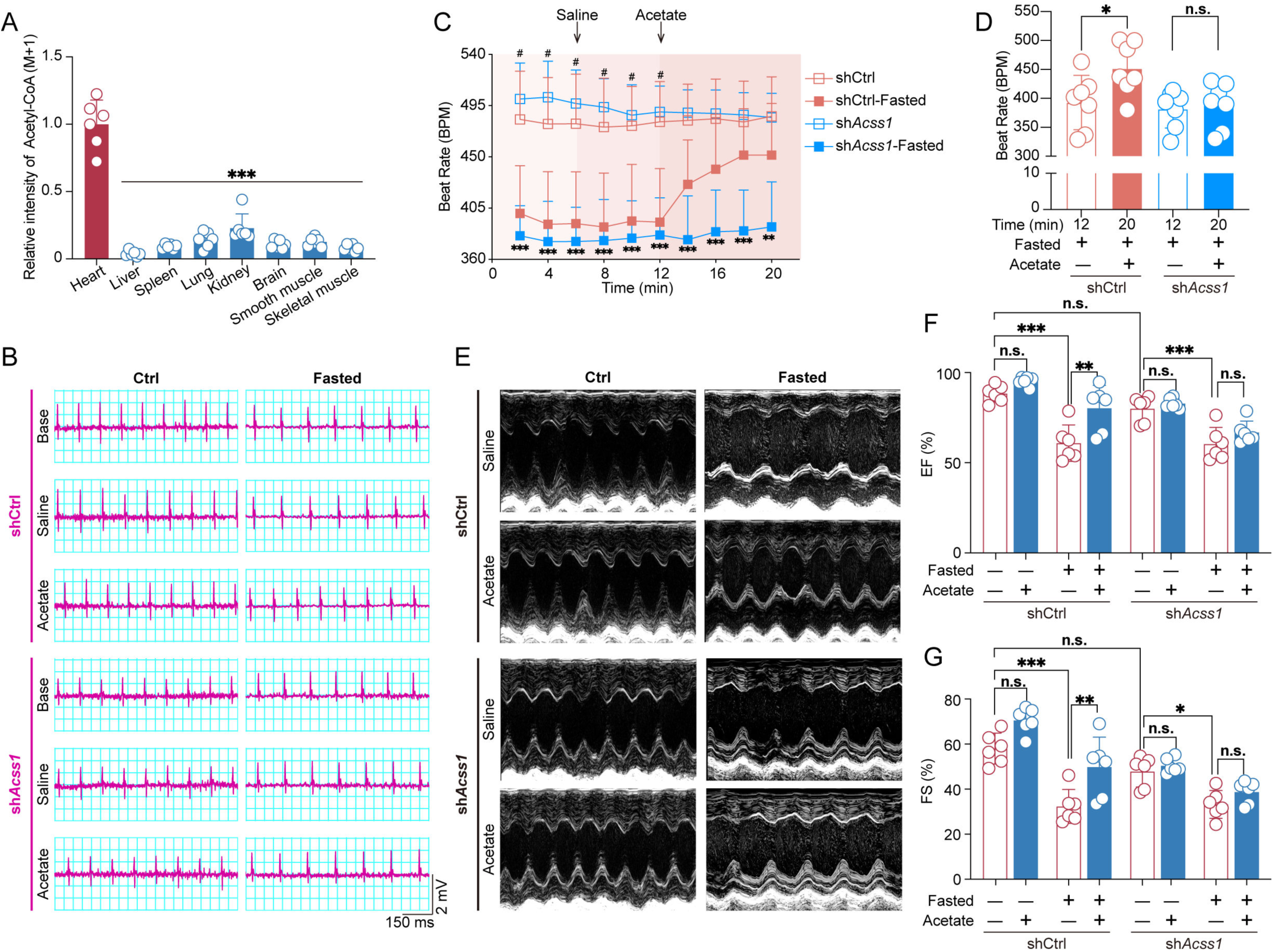
Hepatic Acetate Maintains Cardiac Function via Acss1 *in vivo* under Energy Restriction. (A) Tissue distribution of ^13^C-acetyl-CoA in 18-h-fasted C57BL/6 mice (n = 6) 1 h after intraperitoneal injection of 2-^13^C-acetate (310 mg/kg). Statistical comparisons were performed between each tissue and the heart. (B-D) Representative electrocardiogram (ECG) recordings from 18-h-fasted mice with AAV9-mediated cardiac-specific *Acss1* knockdown at the indicated time points (n = 8). (E-G) Echocardiographic assessment of cardiac function in 18-h-fasted mice with cardiac-specific *Acss1* knockdown, including representative M-mode images (E) and quantitative analysis of ejection fraction (EF, F) and fractional shortening (FS, G) (n = 6). Data are presented as mean ± SD from ≥ 3 biological replicates. Statistical analyses were performed using one-way ANOVA (A), two-way ANOVA (C, D), or three-way ANOVA (F, G), as appropriate. For (A, D, F and G), significance is indicated as \**p* < 0.05, \*\**p* < 0.01, \*\*\**p* < 0.001; n.s., no significant difference. For (B), statistical comparisons between shCtrl and shCtrl-fasted groups are indicated by # (*#p* < 0.05), whereas comparisons between sh*Acss1* and sh*Acss1*-fasted groups are indicated by * (\*\**p* < 0.01, \*\*\**p* < 0.001).

To investigate whether acetate physiologically modulates cardiac function under fasting, we recorded heart rate using Lead II electrocardiography (ECG). Fasting resulted in a marked reduction in heart rate, which was significantly restored by intraperitoneal injection of exogenous acetate (Figure 4B-D). However, in mice with heart-specific Acss1 silencing, acetate supplementation failed to rescue the bradycardia phenotype (Figure 4B-D), indicating a requirement for Acss1 in acetate-mediated chronotropic support. To further evaluate *in vivo* cardiac function, echocardiography was performed in fasted mice. Fasting markedly impaired left ventricular systolic function in both wild-type and cardiac-specific Acss1 knockdown mice. This impairment was significantly reversed by acetate administration in wild-type mice, but not in Acss1-deficient mice, as indicated by increased ejection fraction (EF) and fractional shortening (FS), reflecting improved global systolic performance and myocardial contractility, respectively. In contrast, stroke volume (SV) and cardiac output (CO), which represent beat-to-beat ejection volume and overall cardiac pumping capacity, respectively, remained largely unchanged across conditions (Figure 4E-G and S6B-C). Together, our findings establish that circulating acetate originating from fatty acids oxidation in liver is a critical fuel for maintaining heart rate and cardiac function during fasting in a Acss1-dependent manner.

## DISCUSSION

Our results demonstrate that, under energy-deprived conditions, hepatic production of acetate plays a critical role in sustaining cardiac electrical activity and contractile function. Cardiomyocytes utilize acetate via Acss1, enabling sustained ATP production and electrical activity. This metabolic flexibility highlights the liver-heart interaction in conferring heart ability to adapt to systemic energy scarcity. It is necessary for us to deeply understand the possible reasons why acetate is used by cardiac cells as a crucial fuel in energy-deficient states. Based on collective analyses of our results and other literatures, we suggest that the following aspects should be taken into account.

Firstly, upon fasting, serum acetate can be sustained at the levels as high as 454.2 ± 92.8 μM to sufficiently maintain cardiac electrical activity and contractile function. The physiological acetate levels in normal mice oscillate within a range of 128.9-229.3 μM. The median serum acetate levels show an approximate two-fold increase in fasted mice compared to normally fed mice (Figure 1A), leading to acetate levels in fasted mice being much higher than 250 μM, a near-physiological concentration at which acetate alone could sustain stable cardiac beating and preserve conduction dispersion in the *ex vivo* Langendorff model (Figure 1F-I). These observations demonstrate that upon fasting the serum acetate could be sustained at the levels for efficiently nourishing cardiac tissues. Although propanoate, ketone bodies (3-HB and AcAc) and long-chain fatty acids are also accumulated in serum (Figure 1B-E and S1A-B), none of them alone could support continuous *ex vivo* myocardial contractions with preserved conduction dispersion and normal sinus rhythm (Figure S2A-C). In contrast, lactate levels were reduced in fasted mice to concentrations (1.20 ± 0.22 mM) insufficient to maintain cardiac electrical activity and contractile function *ex vivo* (Figure S1D), despite the ability of supraphysiological concentrations (10 mM) to support relatively normal beating (Figure S2G-I). Similarly, pyruvate sustained cardiac activity at 10 mM but failed to do so at 1 mM (Figure S2J-L). Notably, physiological serum pyruvate levels (0.12 ± 0.01 mM) ^16^ fall well below this functional threshold, despite remaining unchanged during fasting (Figure S1E). Butyrate showed a similar pattern, supporting cardiac activity at 10 mM but not at 5 mM (Figure S2D-F). However, circulating butyrate levels in mice are typically ∼100 μM ^17^, far below the effective range observed *ex vivo*. Likewise, reducing glucose to 1 mM impaired cardiac electrical activity and contractile function, whereas acetate supplementation restored sustained myocardial contraction under reduced glucose availability (Figure 1J-K and S3). In this context, it is reasonable and spontaneous that acetate stands out serving as an emergency fuel for heart.

Secondly, acetate can be selectively consumed by cardiomyocytes, and shows the highest preference in comparation with other short-chain fatty acids and ketone bodies. In an assessment of which kind of murine cell being able to consume acetate, several kinds of primary murine cell lines including cardiomyocytes, C2C12 myoblasts, differentiated C2C12 myotubes (D-C2C12), primary skeletal muscle cells (MSMCs), primary vascular smooth muscle cells (MVSMCs), cardiac fibroblasts (CFs), primary renal proximal tubular epithelial cells (TECs) and HT22 neuronal cells were cultured in 2-^13^C-acetate-containing medium for 24 h, followed by determination of 2-^13^C-acetate consumption. Interestingly, among the tested cell lines, only cardiomyocytes exhibited considerable ^13^C-acetate consumption under glucose-deprived conditions, indicating the uniqueness of cardiomyocyte in consumption of acetate (Figure 2A). This conclusion is confirmed by the results of *in vivo* isotopic tracing assays where 2-^13^C-acetate was administered in fasted mice and detected for ^13^C-acetyl-CoA abundance in various organs. The results indicate that heart exhibits the far more powerful efficiency in consuming acetate than other organs examined (Figure 4A). Furthermore, we examined the preference of cardiomyocyte in consuming short-chain fatty acids and ketone bodies. As shown in Figure 2B, it is acetate, rather than other short-chain fatty acids and ketone bodies, that was utilized by cardiomyocytes with the highest preference when glucose was absent.

Thirdly, acetate may be catabolized more readily than other short-chain fatty acids and ketone bodies. As a high energy-consuming organ, heart needs to extract energy with high efficiency, and spontaneously favors fuels readily degraded, in particular, in energy emergency states such as fasting. Our data indicate that the mitochondria- localized Acss1 shows a selectively high expression in mouse cardiac tissue (Figure 3A), and play the key role in cardiomyocytes consumption of acetate (Figure 3B). Biochemically, acetate can be directly converted to acetyl-CoA by Acss1, clearly more convenient than AcAc and 3-HB which need 2 and 3 enzyme-catalyzed steps to be converted to acetyl-CoA, individually, as well as butyrate and propanoate which need far more steps than acetate in conversion into acetyl-CoA and entry into TCA cycle. In this regard, acetate is the fuel with highest efficiency for heart in energy deficiency states.

In fact, it is reported by Feola K et al that hepatic ketogenesis is not required for starvation adaptation in mice. In this study, mouse was knocked out for 3-hydroxy-3-methylglutaryl-CoA synthase 2 (HMGCS2), the rate-limiting enzyme in the formation of ketone bodies, and observed for starvation adaptation. HMGCS2 knockout-caused hepatic ketogenic deficiency failed to result in any defects in starvation adaptation ^18^. Consistent with our finding, HMGCS2 deficient mice exhibited higher levels of plasma acetate in response to fasting. Coincidentally, the study by Yeh CY et al also reveals that ketogenesis is dispensable for the metabolic adaptations to caloric restriction employing HMGCS2 deletion mice ^19^. Both studies suggest that acetate may play an important role in the adaptions of mouse to starvation or caloric restriction by fueling extrahepatic organs. These studies indirectly support our finding that acetate is an emerging fuel for heart in energy deficiency.

To summarize, in this study we establish acetate which is produced in liver from free fatty acids as a cardiac energy source in energy deficient conditions. Our finding not only provide an unexpected metabolic circuit that directly links hepatic energy mobilization to cardiac demand in energy-deficient state, but also shed a light on the therapeutic potential of acetate for energy-deficient cardiac disorders such as heart failure and so on.

## Supporting information

Supplemental Movie for Figure2D FigureS4A FigureS5A

## RESOURCE AVAILABILITY

## Lead contact

Further information and requests for resources and reagents should be directed to and will be managed by the lead contact, Qinxi Li.

## Materials availability

This study did not generate new unique reagents.

## Data and code availability

The detailed raw data generated in this study have been deposited in the Mendeley Data: https://data.mendeley.com/datasets/xzrvwtdprx/2. This paper does not report original code. Any additional information required to reanalyze the data reported in this paper is available from the lead contact upon request.

## ACKNOWLEDGMENTS

We thank members of the Qinxi Li’s laboratory for productive discussions and comments on this manuscript.

## AUTHOR CONTRIBUTIONS

J.W. and Q.L. conceived the project. J.W. and Yiran Zhang designed most experiments. J.W., T.R., Yiran Zhang, L.Y., H.Y., Z.W., Ying Zhang, Songyang Li and Shitao Liang performed experiments. T.R., J.L., B.J., J.H., G.L. and Q.L. helped with data analysis and discussion. Q.L. and J.W. wrote the manuscript. All authors commented on the manuscript.

## STAR★METHODS

Detailed methods are provided in the online version of this paper and include the following:

## KEY RESOURCES TABLE

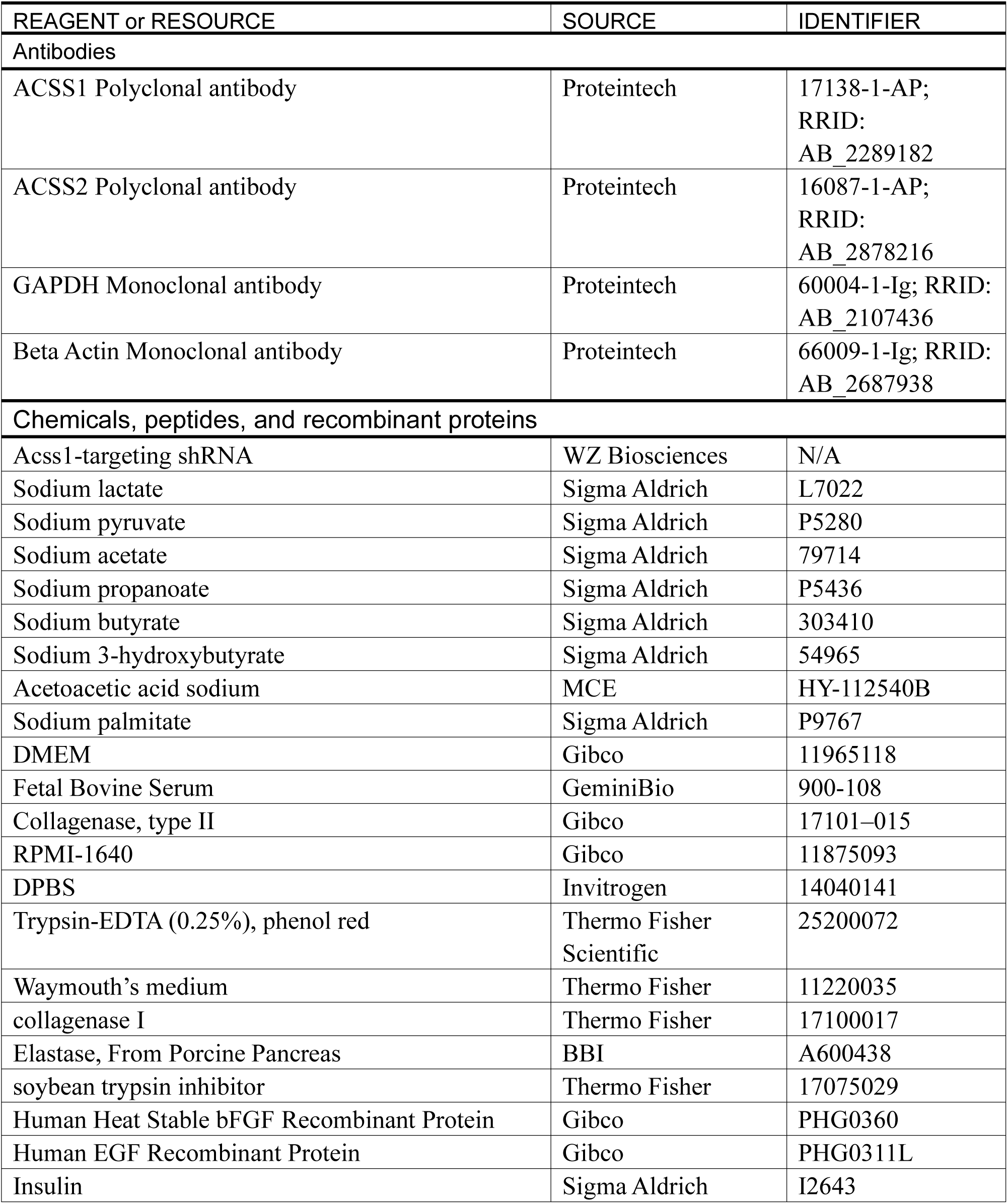
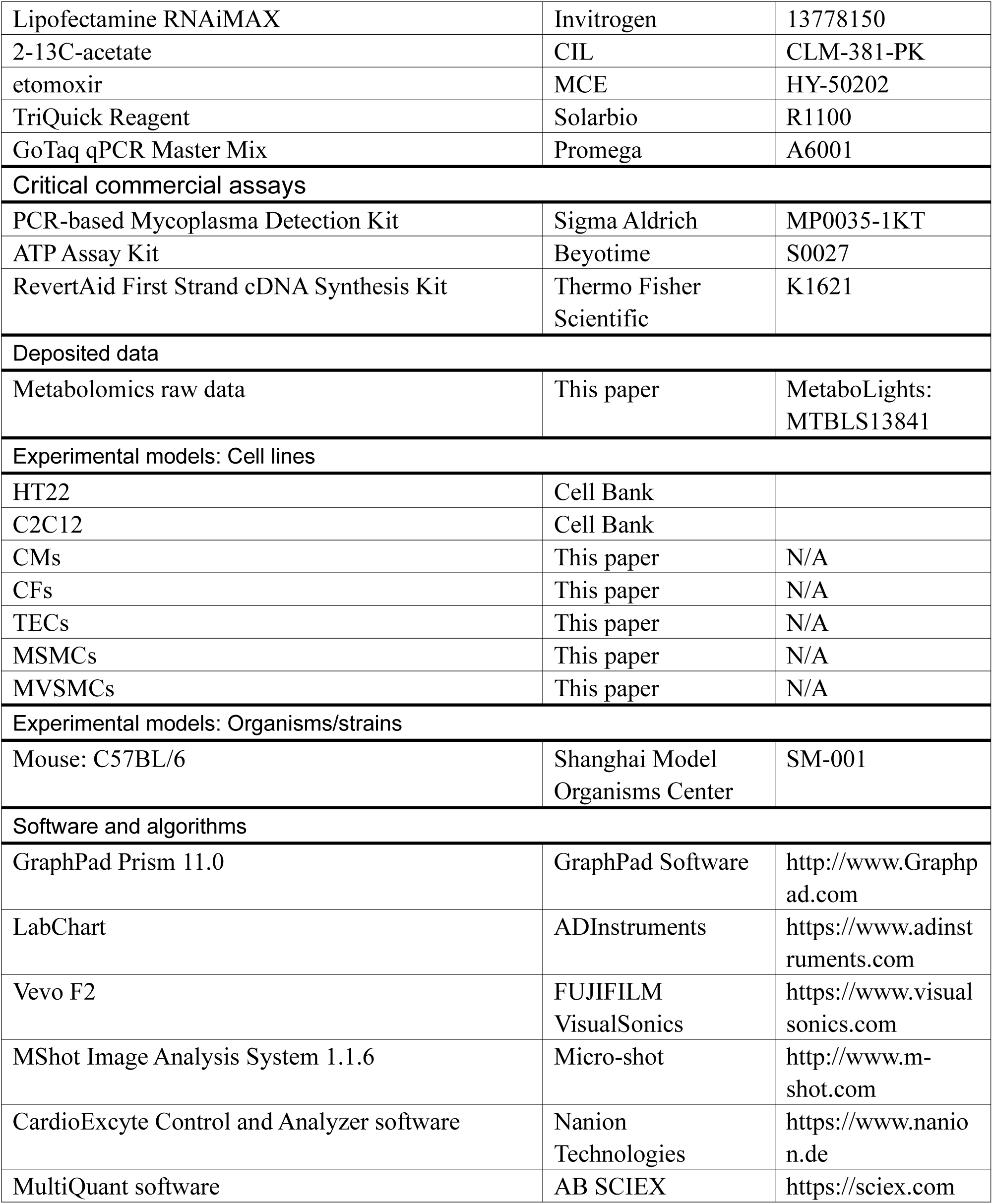

## EXPERIMENTAL MODEL AND STUDY PARTICIPANT DETAILS

### Mice

All animal experiments were approved by the Animal Ethics Committee of Xiamen University (Approval No. XMULAC20240210) and conducted in accordance with institutional and national guidelines. Male C57BL/6 mice (6-8 weeks old) were housed under specific pathogen-free (SPF) conditions with a 12-hour light/dark cycle and ad libitum access to food and water. Cardiomyocyte-specific Acss1 knockdown was achieved by tail-vein injection of AAV9 encoding an Acss1-targeting shRNA (5 × 10^11^ vg per mouse; WZ Biosciences) under a cardiomyocyte-specific promoter; a non-targeting shRNA served as control. The target sequences of shRNA for mouse Acss1: GCATGGCAAGGACCATCTATG. Knockdown efficiency was verified in cardiac tissue by qPCR three weeks post-injection prior to functional analyses. Blood glucose was measured from tail-vein blood using a glucometer (Roche). Blood acetate and lactate was measured from tail-vein blood using NMR. Serum for NMR and MS analyses was obtained by retro-orbital blood collection followed by coagulation and centrifugation. Heart tissues were harvested for Western blotting, and mice were subjected to ECG recording, echocardiography, and *ex vivo* Langendorff perfusion as indicated.

### Isolation of Several Types of Primary Murine Cells

#### Primary Cardiomyocytes (CMs) and Cardiac Fibroblasts (CFs) Isolation

Primary cardiomyocytes were isolated from the hearts of neonatal mice (postnatal day 1-3) using a sequential enzymatic digestion protocol. Briefly, neonatal hearts were excised, rinsed in cold phosphate-buffered saline (PBS), and minced into small fragments. Tissues were then subjected to enzymatic dissociation in 0.1% trypsin (in PBS) at 4 °C overnight (8-12 h). The digestion was neutralized with an equal volume of DMEM (11965118, Gibco) supplemented with 10% fetal bovine serum (FBS, 900-108, Gemini). The cell suspension was centrifuged at 1,000 rpm for 5 min and washed twice with FBS-free DMEM. This was followed by further digestion using 0.1% collagenase type II (17101–015, Gibco) at 37 °C for 3-5 cycles. The resulting suspension was filtered through a 70 μm cell strainer to remove undigested tissue. To separate fibroblasts from cardiomyocytes, cells were pre-plated in culture dishes for 2 hours. Adherent cardiac fibroblasts were retained and expanded for subsequent experiments (passages 2-4). Non-adherent cells, enriched for cardiomyocytes, were collected and cultured in fresh medium. After 48 hours, when cardiomyocytes had fully attached, the culture medium was replaced. The spontaneous beating of cultured cardiomyocytes was monitored and recorded using an inverted microscope equipped with a time-lapse video recording system (MShot Image Analysis System 1.1.6).

#### Primary Mouse Renal Proximal Tubular Epithelial Cells (TECs) Isolation

Male C57BL/6 mice (2-3 weeks old) were euthanized by cervical dislocation. Kidneys were aseptically excised, rinsed in ice-cold RPMI-1640 (11875093, Gibco), and decapsulated. Cortical tissue was dissected, minced in cold RPMI-1640, and centrifuged at 1500 rpm for 5 min. The pellet was digested in 5 mL of collagenase II at 37 °C for 30 min with gentle agitation. Following digestion, the suspension was sequentially filtered through 250 μm and 80 μm strainers. Enzymatic activity was quenched with pre-warmed neutralization buffer. Cells were pelleted, resuspended in pre-warmed TECs culture medium, and seeded into T75 flasks. Cultures were maintained at 37 °C in 5% CO₂, with the first medium change at 48 h and every other day thereafter. Cells were used after reaching confluence.

#### Primary Mouse Skeletal Muscle Cells (MSMCs) Isolation

Skeletal muscles were harvested from the hind limbs of male C57BL/6 mice (6-8 weeks old), minced, and digested in 0.2% type II collagenase in calcium-containing DPBS (14040141, Invitrogen) at 37 °C for 30 minutes. After centrifugation (1800 g, 5 min), the pellet was further digested with 0.25% trypsin-EDTA (1 mM EDTA; AM9260G, Invitrogen) for 10 minutes at 37 °C. The resulting suspension was passed through a 40 μm cell strainer, resuspended in DMEM containing 20% FBS and 1% penicillin-streptomycin, and plated in fresh medium.

#### Primary Mouse Vascular Smooth Muscle Cells (MVSMCs) Isolation

Thoracic and abdominal aortas were isolated from anesthetized male C57BL/6 mice (6-8 weeks old) following isoflurane inhalation. After dissection, aortic segments (excluding heart and carotid arteries) were rinsed with 70% ethanol and washed in Ca²⁺/Mg²⁺-containing DPBS, then transferred to Waymouth’s medium (11220035, Thermo Fisher) supplemented with 1% antibiotic-antimycotic, 1% nonessential amino acids, 1% HEPES, 1% L-glutamine, and 3% sodium bicarbonate. Aortic tissues were minced and digested in smooth muscle growth medium containing 20% FBS, 1 mg/mL collagenase I (17100017, Thermo Fisher), 0.19 mg/mL elastase (39445-21-1, BBI), 0.25 mg/mL soybean trypsin inhibitor (17075029, Thermo Fisher), and growth factor supplements (FGF, PHG0360, Gibco; EGF, PHG0311L, Gibco; insulin, I2643, Sigma-Aldrich), incubated at 37 °C with 5% CO₂ for 3-4 h. Following digestion, the suspension was centrifuged at 200 g for 5 min, and the pellet was resuspended in smooth muscle growth medium. Cells were plated without medium change for 48-96 h to promote primary MVSMCs outgrowth.

All primary murine cells were maintained in DMEM supplemented with 10% FBS, 100 U/mL penicillin, and 100 μg/mL streptomycin at 37 °C in a humidified atmosphere with 5% CO₂.

### Cell Culture, Transfections and Cell Treatments

HT22 and C2C12 cells were purchased from the Cell Bank of the Chinese Academy of Sciences (Shanghai, China) and verified by the supplier. All cell lines were confirmed to be free of mycoplasma contamination using a PCR-based Mycoplasma Detection Kit (MP0035-1KT, Sigma-Aldrich). Cells were cultured in Dulbecco’s Modified Eagle Medium supplemented with 10% fetal bovine serum at 37 °C in a humidified incubator with 5% CO₂. Myogenic differentiation of C2C12 cells was initiated upon reaching 80-90% confluency. Cells were cultured in differentiation medium consisting of high- glucose DMEM supplemented with 2% horse serum and 1% penicillin-streptomycin for 7 consecutive days, with complete medium replacement every 48 hours.

Small interfering RNAs (siRNAs) targeting Acss1 (siRNA #1: GGCTAAAGATCAACCAGTT; siRNA #2: GGACACTCCTTACCATACT; siRNA #3: CCAATACACTGAAGAGACA) and a non-targeting control siRNA were purchased from Thermo Fisher Scientific. For RNAi experiments, cells were transfected twice with 100 nM siRNA using Lipofectamine RNAiMAX (13778150, Invitrogen) according to the manufacturer’s protocol. Knockdown efficiency was evaluated by Western blot.

For all experimental treatments, cells were seeded into 35 mm dishes and pre-incubated for 24 hours. To assess metabolite utilization, cells were washed with PBS and incubated in glucose-free DMEM (1 mL) supplemented with 10% FBS and one of the following substrates for 24 hours: 2-^13^C-acetate (CLM-381-PK, CIL), pyruvate, lactate, propionate, butyrate, or 3-hydroxybutyrate. To simulate energy-restricted conditions—limiting both glucose and fatty acid availability—cells were incubated in glucose-free DMEM (1 mL) supplemented with 10% FBS and etomoxir (40 μM; HY-50202, MCE) for 1-2 hours. The culture media from treated cells were collected for subsequent nuclear magnetic resonance (NMR) analysis.

## METHOD DETAILS

### Electrocardiography (ECG)

Electrocardiograms (ECGs) were recorded as previously described ^20^. Mice were lightly anesthetized with 1% isoflurane and maintained on a temperature-controlled platform. Lead II ECG signals were acquired using electrodes placed on the plantar surfaces of the limbs, with the right forelimb as the negative electrode, the left hindlimb as the positive electrode, and the right hindlimb as the ground. Signals were recorded using a PowerLab 15T four-channel physiological recording system (ADInstruments).

### Echocardiography

Transthoracic echocardiography was performed using a high-resolution ultrasound system (Vevo F2, VisualSonics) as previously described ^21^. Mice were anesthetized with inhaled isoflurane and maintained on a heated platform at 37 °C, with heart rate stabilized at approximately 400 beats per minute. After chest hair removal and application of ultrasound coupling gel, cardiac function was assessed using a 40-MHz transducer. M-mode images were acquired from parasternal long-axis view, and measurements were averaged over three consecutive cardiac cycles. Left ventricular end-diastolic and end-systolic diameters (LVEDD and LVESD), heart rate (HR), ejection fraction (EF), fractional shortening (FS), stroke volume (SV), and cardiac output (CO) were quantified using the manufacturer’s analysis software.

### Langendorff Hearts

Langendorff perfusion was performed according to a previously established protocol ^11^. Male C57BL/6J mice (10-12 weeks old) were injected intraperitoneally with 500 U of heparin to prevent coagulation. After 10 minutes, mice were euthanized by cervical dislocation, and hearts were rapidly excised in ice-cold Krebs-Henseleit (KH) buffer. The aorta was cannulated and connected to a Langendorff perfusion system. Retrograde perfusion was conducted at 37 °C using oxygenated KH buffer containing: glucose, NaCl, NaHCO₃, KCl, KH₂PO₄, MgCl₂, and CaCl₂; the buffer was continuously bubbled with 95% O₂ and 5% CO₂ to maintain pH at 7.4. For *ex vivo* electrophysiological measurements, a 64-channel epicardial multielectrode array was gently attached to the surface of the left ventricle. Electrical signals were continuously recorded to assess cardiac conduction properties. To evaluate substrate-specific effects on cardiac electrophysiology, glucose in the KH buffer was individually replaced with one of the following metabolites: Sodium lactate (L7022, Sigma-Aldrich), Sodium pyruvate (P5280, Sigma-Aldrich), Sodium acetate (79714, Sigma-Aldrich), Sodium propanoate (P5436, Sigma-Aldrich), Sodium butyrate (303410, Sigma-Aldrich), Sodium 3-hydroxybutyrate (54965, Sigma-Aldrich), Acetoacetic acid sodium (HY-112540B, MCE), or Sodium palmitate (P9767, Sigma-Aldrich).

### Measurement of Intracellular ATP Levels

Intracellular ATP levels were measured using an ATP Assay Kit (S0027, Beyotime) according to the manufacturer’s instructions. Briefly, cells were seeded into 6-well plates and incubated in glucose-free DMEM (1 mL) supplemented with 10% FBS and etomoxir (40 μM) for 6 hours after stabilization. Subsequently, 200 μL of ATP lysis buffer was added to each well to lyse the cells. The lysates were centrifuged at 12,700 rpm for 10 minutes at 4 °C, and the supernatants were collected. ATP concentrations were quantified by mixing the supernatants with the ATP detection reagent and measuring luminescence using a multimode microplate reader.

### CardioExcyte 96 Assay

Electrical activity of cardiomyocytes was monitored using the CardioExcyte 96 system in impedance (IMP) mode ^22^. Cardiomyocytes were seeded as confluent monolayers on 96-well sensor plates with integrated electrodes to ensure stable signal acquisition. Synchronized contractile activity induced impedance fluctuations resulting from ion flux–mediated potential changes within the narrow electrode–cell cleft. Owing to the high seal resistance and large electrode area, impedance signals reflected the averaged electrophysiological and contractile behavior of the cardiomyocyte population in each well. Data were acquired and analyzed using CardioExcyte Control and Analyzer software (Nanion Technologies) to quantify beat rate and contractile parameters.

### Western blot

Cells or tissues were lysed in ice-cold buffer containing Tris-HCl, NaCl, EDTA, phosphatase inhibitors, and 1% Triton X-100, supplemented with protease inhibitors. Lysates were sonicated and clarified by centrifugation (12,700 rpm, 10 min, 4 °C). Tissue samples were homogenized to ensure complete lysis. Equal amounts of protein were resolved by SDS-PAGE and transferred to PVDF membranes (Roche). Membranes were blocked with 5% BSA in TBST and incubated with primary antibodies followed by HRP-conjugated secondary antibodies. Proteins were visualized by enhanced chemiluminescence. Primary antibodies against ACSS1 (17138-1-AP), ACSS2 (16087-1-AP), GAPDH (60004-1-Ig), and Actin (66009-1-Ig) were obtained from Proteintech.

### qPCR

Total RNA from heart tissue was extracted using TriQuick Reagent (R1100, Solarbio). cDNA was synthesized from 1 μg of RNA using the RevertAid First Strand cDNA Synthesis Kit (K1621, Thermo Fisher Scientific). Quantitative PCR was performed with GoTaq qPCR Master Mix (A6001, Promega). Relative mRNA levels were normalized to Actin. Primer sequences were as follows: Actin, 5′-AGGGAAATCGTGCGTGACAT (forward) and 5′-CGTTGCCAATAGTGATGACC (reverse); Acss1, 5′-GCAGGCTATCTACTGTATGCCG (forward) and 5′-AGGACTGTGGTAGCTCCATTGC (reverse).

### Measurement of Tissue Acetyl-CoA Levels by LC-MS

Tissue acetyl-CoA levels were measured by liquid chromatography-mass spectrometry (LC-MS) to assess acetate utilization across different organs in mice fasted for 18 hours with free access to water (10-12 weeks old, male), as previously described ^23^. For metabolite extraction, equal amounts of tissue from various organs were homogenized in ice-cold methanol/water (4:1, v/v). Samples were centrifuged at 12,000 × g for 20 minutes at 4 °C, and the supernatants were transferred to new Eppendorf tubes and dried at 4 °C. Pellets were then resuspended in acetonitrile/water (1:1, v/v) and transferred to HPLC vials for LC-MS analysis. Metabolite separation was performed using a SCIEX ExionLC AD system equipped with a Millipore ZIC-pHILIC column (5 μm, 2.1 × 100 mm, PN: 1.50462.0001) maintained at 40 °C. The mobile phase consisted of solvent A (15 mM ammonium acetate and 0.3% ammonium hydroxide in LC-MS-grade water) and solvent B (90% acetonitrile in LC-MS-grade water). The flow rate was 0.2 mL/min under the following gradient conditions: 95% B for 2 minutes, linearly decreased to 45% B over 13 minutes, held at 45% B for 3 minutes, then returned to 95% B and maintained for 4 minutes for re-equilibration. Mass spectrometry was conducted using a SCIEX QTRAP system equipped with a Turbo V™ electrospray ionization (ESI) source operating in negative ion mode. The source parameters were as follows: ion spray voltage, -4,500 V; Gas 1, 40 psi; Gas 2, 50 psi; curtain gas, 35 psi. Metabolites were detected in multiple reaction monitoring (MRM) mode. Data acquisition and analysis were performed using MultiQuant software (AB SCIEX), and metabolite levels were normalized to an internal standard.

### Nuclear Magnetic Resonance (NMR) Spectroscopy

For nuclear magnetic resonance (NMR) analysis, cell culture medium samples were collected after treatment and centrifuged at 3,000 rpm for 5 minutes at 4 °C. A total of 450 μL of the supernatant was transferred into standard 5 mm NMR tubes. For preparation of mice serum sample, 100 μL serum was mixed with 400 μL deuterium oxide (D₂O), and centrifuged (3000 g) at 4 °C for 10 min. The supernatants (400 μL) were transferred into 5 mm NMR tubes for NMR measurement. A coaxial inner tube containing 200 μL of D₂O with 1 mM sodium 3-(trimethylsilyl) propionate-2,2,3,3-d₄ (TSP) was inserted to serve as both a chemical shift reference (δ 0.00 ppm) and an internal standard for metabolite quantification.

NMR spectra were acquired on two high-field NMR spectrometers: a Bruker Avance III 850 MHz equipped with a TCI cryoprobe (Bruker BioSpin, Germany), provided by the College of Chemistry and Chemical Engineering at Xiamen University, and a Bruker Avance III 600 MHz spectrometer housed in the Core Facility of Biomedical Sciences at Xiamen University. One-dimensional (1D) Carr-Purcell-Meiboom-Gill (CPMG) pulse sequences were used to acquire spectra from culture medium and serum samples with water suppression. The pulse sequence [RD-90°-(τ-180°-τ)ⁿ-ACQ] was applied to attenuate broad signals from macromolecules, thereby enhancing the resolution and detection of low-molecular-weight metabolites.

### Metabolomics Analysis

For serum metabolomics analysis, male C57BL/6 mice (10-12 weeks old) were fasted for 18 hours with free access to water. Blood samples were collected and immediately placed on ice. Serum was isolated by centrifugation at 3,000 × g for 10 minutes at 4 °C and stored at -80 °C until analysis. After thawing and vortexing, 30 μL of serum from each sample was aliquoted into 1.5 mL centrifuge tubes and divided into six portions: four for multi-channel derivatization analysis (30 μL/channel), one as a backup, and one for pooled sample preparation. For pooled sample generation, 75 μL of serum from each sample was combined, vortexed thoroughly, and labeled as a reference sample. For extraction, 90 μL of pre-chilled LC-MS grade methanol was added to each tube containing 30 μL of serum. Samples were vortexed, placed at -20 °C for 1 hour to precipitate proteins, and then centrifuged at 12,000 rpm for 10 minutes at 4 °C. A 90 μL aliquot of the supernatant was transferred to a new centrifuge tube and dried under a gentle stream of nitrogen.

Four-channel chemical derivatization (targeting amines/phenols, carboxyls, hydroxyls, and carbonyls) was performed according to standard operating procedures prior to LC-MS analysis. Chromatographic separation was conducted using an Agilent Eclipse Plus C18 column (150 mm × 2.1 mm, 1.8 μm) on an Agilent 1290 Infinity LC system coupled to an Agilent 6546 Q-TOF mass spectrometer. The mobile phases consisted of 0.1% formic acid in water (A) and 0.1% formic acid in acetonitrile (B). The gradient program was as follows: 25% B at 0 minutes, increased linearly to 99% B over 10 minutes, held at 99% B for 5 minutes, returned to 25% B at 15.1 minutes, and maintained until 18 minutes. The column temperature was set to 40 °C, and the flow rate was 400 μL/min. The MS ion source parameters were: gas temperature, 325 °C; dry gas, 8 L/min; nebulizer, 35 psi; sheath gas temperature, 400 °C; sheath gas flow, 12 L/min; scan range, m/z 220-1000.

Metabolites were annotated by matching MS data against the HMDB (http://www.hmdb.ca) and METLIN (https://metlin.scripps.edu) databases. Differentially abundant metabolites (DAMs) were subjected to metabolic pathway enrichment analysis based on the KEGG database using MetaboAnalyst 6.0 (https://www.metaboanalyst.ca/). Raw metabolomics data have been deposited in the MetaboLights repository (https://www.ebi.ac.uk/metabolights) under accession number MTBLS13841. Although the dataset is currently under review and not publicly available, it can be accessed for peer review via the following private link: https://www.ebi.ac.uk/metabolights/MTBLS13841.

## QUANTIFICATION AND STATISTICAL ANALYSIS

All statistical analyses were performed using GraphPad Prism (version 11.0) and Microsoft Excel. Data are presented as mean ± standard deviation (SD) unless otherwise indicated. For comparisons between two groups, a two-tailed unpaired Student’s *t*-test was used. For experiments involving more than two groups, one-way, two-way or three-way analysis of variance (ANOVA) was performed, followed by appropriate post hoc multiple-comparison tests as specified in the figure legends. A *P* value < 0.05 was considered statistically significant.

## SUPPLEMENTAL INFORMATION

**Supplementary Video Data.** Representative videos of spontaneous beating of cardiomyocytes under the indicated experimental conditions, corresponding to the traces shown in Figure 2D, Figure S4A, and Figure S5A.

---

**Figure S1.**
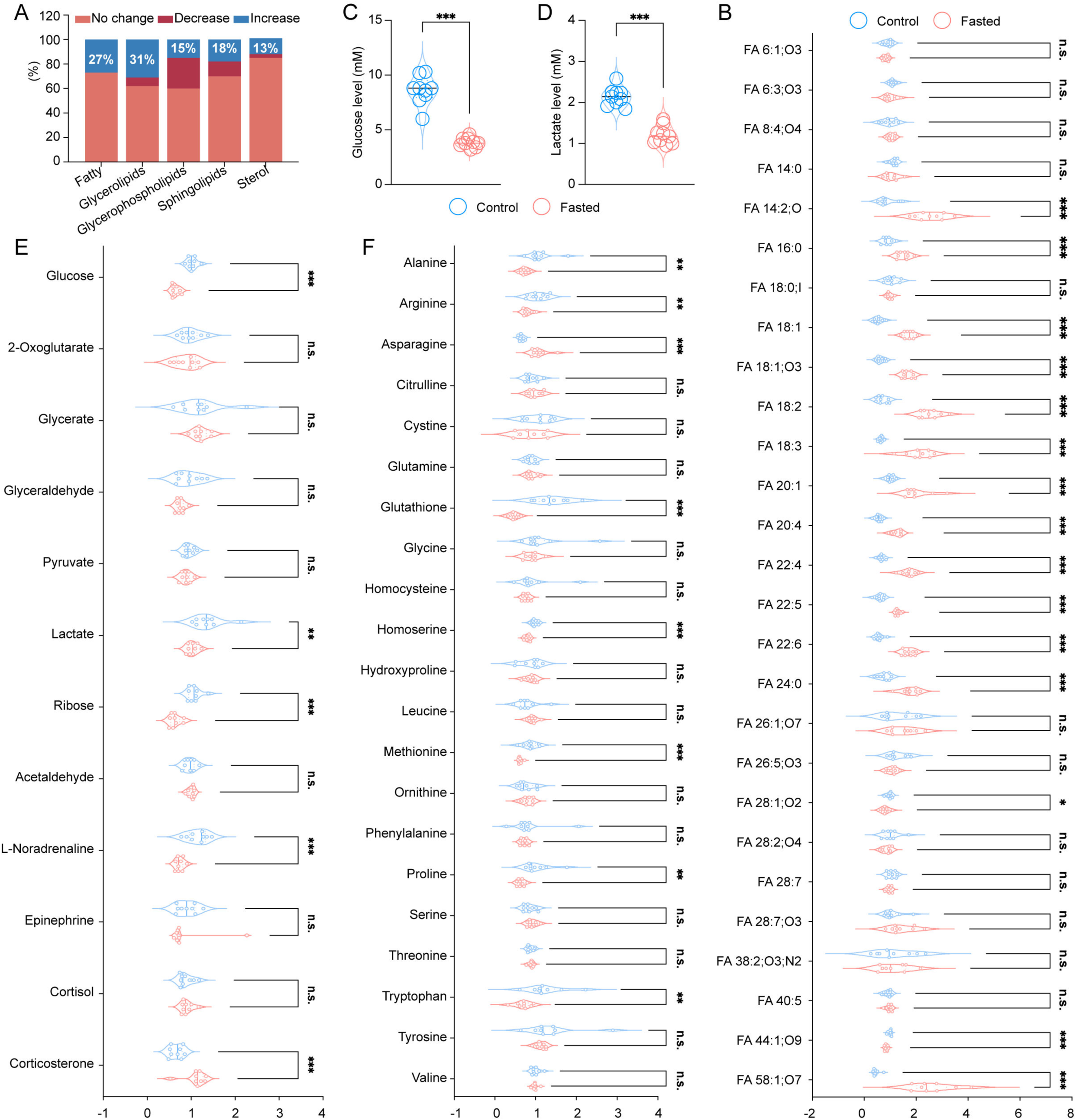
Fasting induced marked metabolic alteration in serum from mice. Relative serum levels of lipid metabolites (A, B), glucose-related metabolites (C-E), and amino acids (F) in mice fasted for 18 h (n = 9). Values are expressed as mean ± standard deviation (SD) and analyzed using two-tailed unpaired Student’s *t*-tests (\**p* < 0.05, \*\**p* < 0.01, \*\*\**p* < 0.001; n.s., no significant difference).

**Figure S2.**
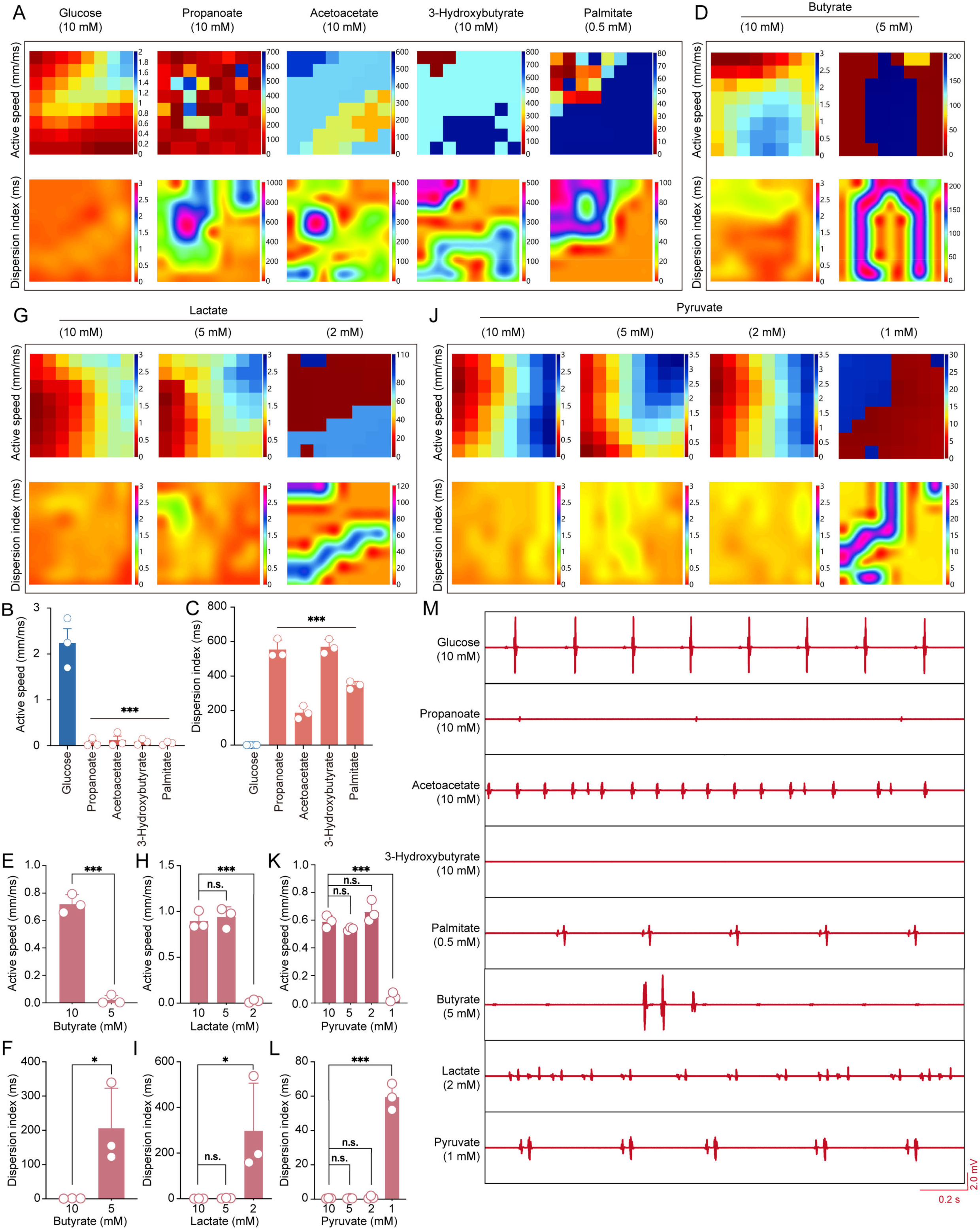
Metabolite-dependent regulation of cardiac electrical activity in Langendorff-perfused hearts. Wild-type mouse hearts were perfused *ex vivo* using the Langendorff system. (A-C) Representative maps of ventricular activation speed and dispersion index (A) and corresponding quantification of activation speed (B) and dispersion index (C) in hearts exposed to the indicated metabolites at 10 mM (n = 3). (D-F) Representative maps (D) and quantification of activation speed (E) and dispersion index (F) in hearts perfused with butyrate at 10 mM and 5 mM (n = 3). (G-I) Representative maps (G) and quantification of activation speed (H) and dispersion index (I) in hearts perfused with lactate at 10, 5, and 2 mM (n = 3). (J-L) Representative maps (J) and quantification of activation speed (K) and dispersion index (L) in hearts perfused with pyruvate at 10, 5, 2, and 1 mM (n = 3). (M) Representative electrocardiogram traces from Langendorff-perfused hearts under the indicated conditions. Values are expressed as mean ± SD and analyzed using one-way ANOVA (\**p* < 0.05, \*\**p* < 0.01, \*\*\**p* < 0.001; n.s., no significant difference).

**Figure S3.**
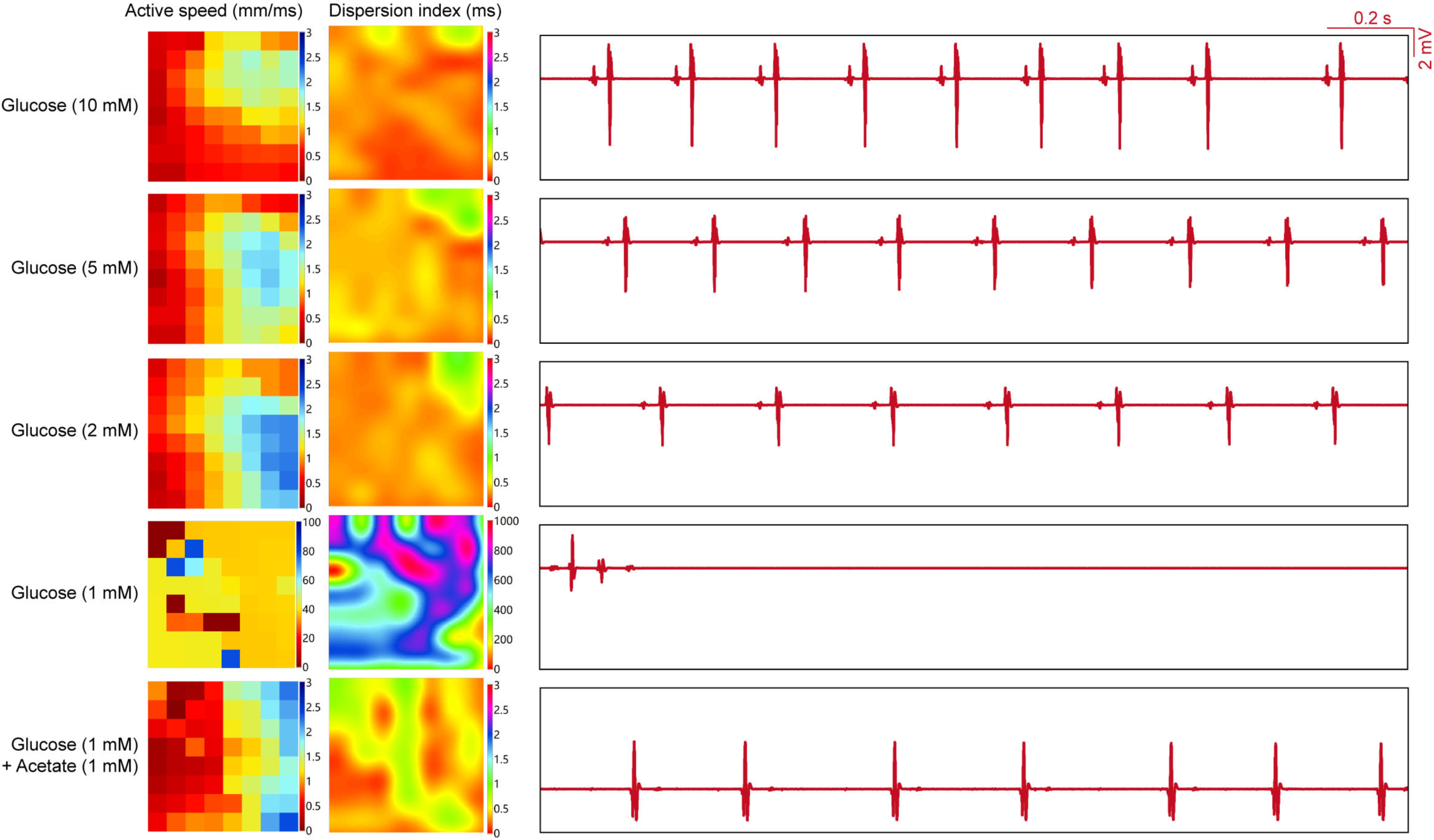
Acetate rescues loss of sustained myocardial contraction under reduced glucose *ex vivo*. Representative ventricular activation speed and dispersion index maps, as well as corresponding electrocardiogram traces, from Langendorff-perfused wild-type mouse hearts exposed to different metabolites (n = 3).

**Figure S4.**
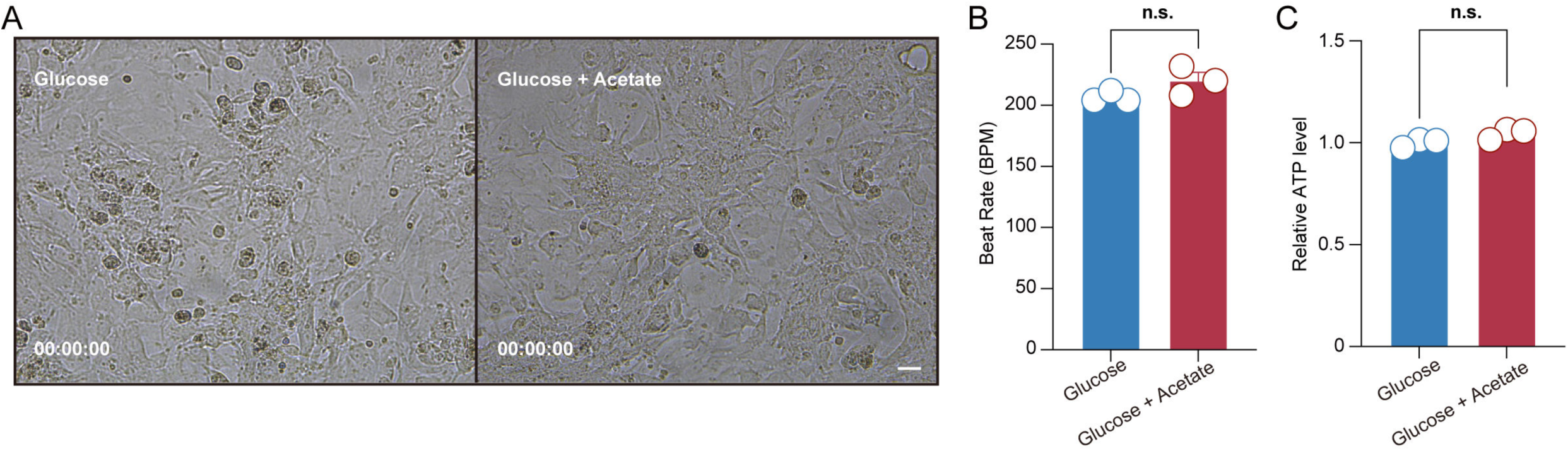
Acetate had minimal effects under nutrient-rich conditions. (A-B) Spontaneous beating of cultured CMs in acetate-supplemented DMEM, monitored (A) and quantified (B) (n = 3). Scale bars represent 20 μm. Video data (A) are provided in the Supplementary Materials. (C) Intracellular ATP levels in CMs after 6 h culture in acetate-supplemented DMEM (n = 3). Values are expressed as mean ± SD and analyzed using two-tailed unpaired Student’s *t*-tests (\**p* < 0.05, \*\**p* < 0.01, \*\*\**p* < 0.001, n.s., no significant difference).

**Figure S5.**
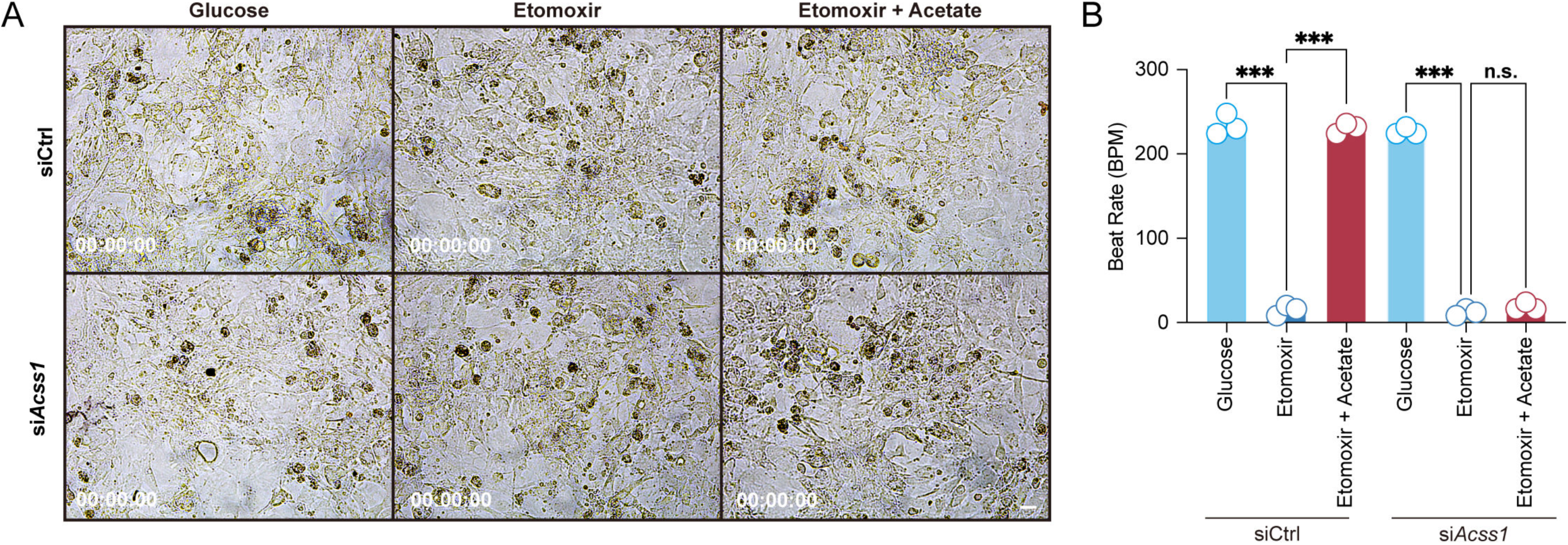
Acss1 loss abolished acetate-mediated CM beating rescue. (A-B) Spontaneous beating of *Acss1*-knockdown CMs cultured for 24 h under energy-restriction conditions (glucose-free medium with 40 μM etomoxir) with acetate supplementation was monitored (A) and quantified (B) (n = 3). Scale bars represent 20 μm. Video data (A) are provided in the Supplementary Materials. Values are expressed as mean ± SD and analyzed using two-way ANOVA (\**p* < 0.05, \*\**p* < 0.01, \*\*\**p* < 0.001, n.s., no significant difference).

**Figure S6.**
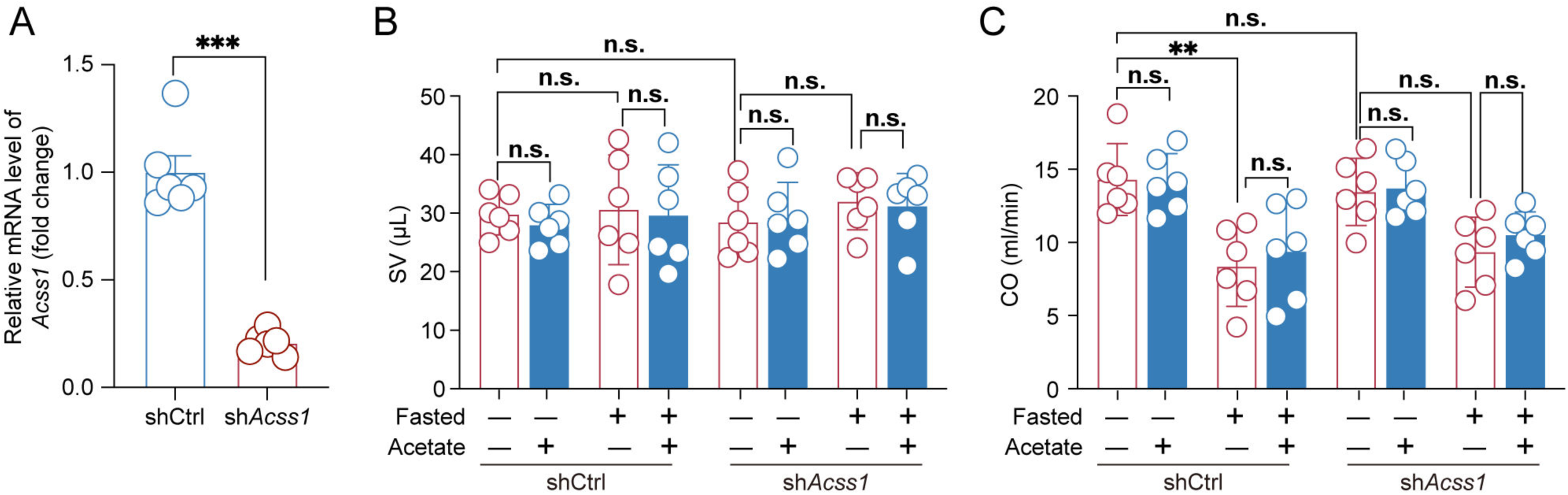
Effects of Acss1 knockdown on cardiac contractile function in mice. (A) Cardiac-specific *Acss1* knockdown in C57BL/6 mice achieved by AAV9-delivered shRNA, validated by qPCR analysis of heart tissue. (B-C) Echocardiographic assessment of cardiac function in 18-h-fasted mice with cardiac-specific *Acss1* knockdown, showing stroke volume (SV) (B) and cardiac output (CO) (C) (n = 6). Values are analyzed using two-tailed unpaired Student’s *t*-tests (A) or three-way ANOVA (B and C) (\**p* < 0.05, \*\**p* < 0.01, \*\*\**p* < 0.001, n.s., no significant difference).

